# Unraveling the impact of hyperleptinemia on female reproduction: Insights from transgenic pig model

**DOI:** 10.1101/2024.05.20.595055

**Authors:** Muhammad Jamal, Yixiao Cheng, Deling Jiao, Wen Cheng, M Di Zou, Xia Wang, Taiyun Wei, Jianxiong Guo, Kaixiang Xu, Heng Zhao, Shaoxia Pu, Chang Yang, Yubo Qing, Baoyu Jia, Honghui Li, Rusong Zhao, Hong-Ye Zhao, Hong-Jiang Wei

## Abstract

**Background:** Infertility is a growing global health concern affecting millions of couples worldwide. Among several factors, an extreme body weight adversely affects reproductive functions. Leptin is a well-known adipokine that serves as an endocrine signal between adiposity and fertility. However, the exact mechanisms underlying the effects of high leptin on female reproduction remain unclear.

**Methods:** Transgenic pig overexpressing leptin (11) were produced by back cross and screened for leptin overexpression, and the growth curve, fat deposition, reproductive performance, apoptosis, serum hormones and cholesterol production, RNA sequencing, and single nucleus RNA sequencing of the leptin-overexpressed pigs and control group were evaluated.

**Results:** Transgenic pig overexpressing leptin (11) were obtained, which exhibited significantly reduced body weight, body size, and back fat thickness. These pigs manifested late onset of puberty (327±48.5 vs 150±6.5 days), irregular estrous behavior characterized by increased inter-estrous interval (28.1±4.2 vs 21.3 ± 0.9 days), and more numbers of mating until pregnancy (at least 3 times). This reproductive impairment in leptin pigs was related to hormonal imbalances characterized by increased levels of FSH, LH, prolactin, E2, P4, and TSH, altered steroidogenesis such as increased levels of serum CE along with steroidogenic markers (STAR, CYP19A), and ovarian dysfunctions manifested by neutrophilic infiltration and low expression of caspase-3 positive cells on leptin pigs ovary. Meanwhile, bulk RNA sequencing of the ovaries also revealed neutrophilic infiltration followed by upregulation of inflammation-related genes. Further, leptin overexpression triggered immune response, suppressed follicle development and luteinization, imposing metabolism dysfunction and hormone imbalance in the ovary by single-nucleus RNA sequencing (snRNA-seq).

**Conclusion:** Low body weight in leptin overexpression pigs adversely affects reproductive performance, causing delayed puberty, irregular estrous cycles, and reduced breeding efficiency. This is linked to metabolic imbalances, increased immune response, and altered ovarian functions. This study provided a theoretical basis for the complex mechanisms underlying leptin, and infertility by employing leptin-overexpressed female pigs.

## Background

Infertility is a complex disorder that affects about 10-15% of couples all over the world. Among several factors, optimum body weight is necessary for initiation and maintenance of reproductive cyclicity [1]. An extreme body weight adversely affects reproductive functions starting from pubertal development, menstrual cyclicity, and pregnancy [2]. Several studies indicated that underweight women had delayed onset of puberty followed by disruption of the menstrual cycle that may lead to prolonged periods of amenorrhea [3–5]. Further, long-lasting underweight also disturbs the gonadotropin secretions that can inhibit female copulatory behavior eventually, leading to infertility [6]. Likewise, overweight females also exhibit a higher incidence of menstrual dysfunction, anovulation, and infertility [7]. This impaired reproduction in both extremes is due to impaired secretions of gonadotropins (FSH, LH) that in turn affect the production of steroids and disrupt ovarian function resulting in infertility [8]. Thus, adequate mass of adipose tissue is required for the onset of puberty and maintenance of fertility in females.

The adipose tissue is regarded as a passive organ for lipids storage. It provides free fatty acids in response to energy demands and works as an endocrine organ that releases several hormones and cytokines [9]. Leptin is an adipocyte-derived hormone that was initially known as a satiety factor controlling feed intake and energy metabolism but later, it was set up to be an essential regulator of several physiological processes including reproduction. In the context of reproduction, leptin induces the gonadotropin-releasing hormone (GnRH) expression, sequentially; peripheral LH and FSH are stimulated to potentiate the follicular growth and steroid production [10]. Shreds of evidence from murine models indicated that leptin deficiency led to infertility, while exogenous administration of recombinant leptin restored fertility. Further, transient elevation of plasma leptin concentrations before adolescence related to changes in neuroendocrine function, it is postulated that the “leptin surge” in childhood may signal the initiation of puberty [11]. In brief, leptin serves as an endocrine signal between the degree of adiposity and fertility, providing a peripheral message to the CNS on the adequacy of nutritional status for reproductive functions.

Genetically engineered mouse models have contributed prodigiously to elucidate the role of leptin in reproduction and it was reported that transgenic mice overexpressing leptin were skinny with accelerated puberty and intact fertility at younger ages followed by successful delivery of healthy pups. However, at older ages, they develop hypothalamic hypogonadism characterized by prolonged menstrual cycles, atrophic ovary, reduced hypothalamic GnRH contents, and poor pituitary luteinizing hormone secretion [7]. Nevertheless, small size, short life, and other differences between humans and rodents limit the ability to model complex diseases like infertility. Contrarily, pigs being large in size and having anatomical, physiological, and genetic similarities are regarded as ideal models for humans. A large number of pig models for cancer, diabetes, cardiovascular, neurodegenerative, and immunological disorders have been developed in recent years [12]. Pig models have been utilized for evaluating reproductive disorders like polycystic syndrome, ovarian cancer, and preeclampsia [13]. Thus, in this study, we utilized the pig model to evaluate the effect of high leptin on female reproduction.

Indeed, transcriptome studies using bulk RNA samples have identified changes in leptin expression in pig ovaries, but interpretation is limited due to the heterogeneity of the organ and the varying effects of leptin on different cell types, which are not captured by bulk sequencing or qPCR. However, recent advances in single-nucleus RNA sequencing (snRNA-seq) enable transcriptomic analysis of large numbers of cells at single-cell resolution with high depth [14]. This method provides insights into the gene expression profiles of individual ovarian cells, allowing for a better understanding of the heterogeneity of the ovarian tissue and the specific effects of leptin on different cell types [15–17]. Based on the leptin-overexpressed pigs already generated in our laboratory [18], in this study, the growth curve, fat deposition, reproductive performance, serum hormones and cholesterol profile, steroidogenesis and apoptosis, and changes at individual ovarian cells of the leptin overexpressed pigs and WT group were evaluated systemically to further elucidate the effect of leptin on female reproduction and its mechanism, and to provide a certain theoretical basis in this area of research.

## METHODS

### Animals

Animals used in this study were regularly maintained in the Laboratory Animal Centre of Yunnan Agricultural University. All animal experiments were performed with the approval of the Animal Care and Use Committee of Yunnan Agricultural University. The female pigs overexpressing leptin were obtained by backcrossing the wild-type (WT) female with already generated leptin overexpressing transgenic boar [18] and offspring were screened for the presence of transgenic vector containing leptin and enhanced green fluorescent protein (EGFP) by polymerase chain reaction (PCR) using primers listed in Table S1, and circulating serum leptin concentrations.

### Polymerase Chain Reaction (PCR)

The genomic DNA from blood of each newborn piglet was extracted using TIANamp genomic DNA kit (Tiangen, Beijing, China) and PCR was carried out using 2×Ex Taq Master Mix (TaKaRa CW0718). The PCR reaction was composed of 0.5 µg of template, 10 µl of Ex Taq Master Mix, 0.4 µM of each primer and distilled water to a final volume of 20 µl. The amplification of each primer pairs was performed in separate reaction with 35 cycles of 94 °C for 30 seconds, 55-60 °C for 30 seconds, and 72 °C for 30 seconds. The PCR products were separated by 1% agarose gel electrophoresis and visualized by staining with DL 2000 DNA Marker (TSINGKE). Depending upon the presence of transgenic vector comprised of leptin gene with EGFP expression, the offspring were divided into two groups: Leptin pigs and WT pigs. Next, the circulating serum leptin concentrations were measured using Porcine Leptin Elisa Kit (SEKP-0278, Solarbio life sciences, China).

### Body Weight and Size Measurements

Body weight (BW) and size including length, height, heart girth, chest depth, chest width, and abdominal circumference of leptin (n=5) and WT (n=4) pigs were measured at 30d intervals from d180∼d390 of age. The live weight of each animal was determined by suspending the animal on an electronic scale balance and the weight of each animal was taken and recorded. The body size measurements were recorded. In brief, body length was measured using the tape rule as the distance from the occipital protuberance to the base of the tail and heart girth was determined by measuring the chest with a tape. Height was measured as a distance from the surface of the ground to the withers using a meter rule and chest depth was measured as the distance from the sternum at the region just in front of the forelimb to the withers, while chest width was recorded as distance between shoulders using meter ruler. Abdominal circumference was measured from the bottom of the flank on one side to the bottom of the flank on the other side of the pig.

### Back Fat Thickness

The back fat thickness of WT and leptin pigs was scanned ultrasonically. Briefly, pigs were immobilized and restrained by the head in a squeeze chute and the image site at the P2 position (at the level of the head of the last left rib) was determined by physical palpation. The pigs were held manually, avoiding any abnormal situation that could stress the animal, and were only scanned in a relaxed posture, permitting accurate measurements. A mix of Eco Gel and isopropyl alcohol was used as a sound-conducting material to allow a better acoustic contact surface between the probe and the skin. An Aplio 500V real-time ultrasound machine (BSM34-0927, Toshiba, Japan) equipped with a 5.0-14.0 MHz linear array transducer (PLT-1005BT, Toshiba) was used for scanning the image site.

### Reproductive Performance

Reproductive parameters comprising of onset of puberty, pattern of estrous expression, and breeding performance were evaluated in leptin and WT pigs. The estrus observation in all pigs was started at 120 days of age and gilts were visually observed two times (9:00 AM and 05:00 PM) a day for the expression of estrus signs in the presence of boar and stages of estrous i.e. proestrus, estrus, met-estrus were ascertained according to previously described estrus scoring system [19] with slight modifications. The age at which gilts displayed first estrus was recorded as the age of puberty and the pattern of estrous expression was recorded until 540 d. The inter-estrous interval was defined as the duration (days) from the start of one estrous to the start of the next estrous. Next, we evaluated the breeding performance of leptin and WT pigs by mating with a fertile male. Briefly, leptin females (n=4) were used for mating, and gilts were mated 2 times in one estrous cycle using the same male. Usually, the first mating was performed 24 hours after standing reflex followed by the second mating 18 hours later. Pregnancy was monitored by non-return to estrous and confirmed by ultrasound scanner for around 33 days after service (HS-101 V, Honda Electronics Co., Ltd., Yamazuka, Japan).

### Serum Hormonal and Lipid Profile

Blood samples were collected at five different stages of the estrous cycle: proestrus, estrus, met-estrus, d6 post-met-estrus, and d12 post-met-estrus for hormonal analysis. Briefly, the pigs were fasted overnight and 5∼10 mL blood was sampled from the jugular vein using a vacutainer (Cat no. 367986, BD Vacutainer SST™, USA) and transported to the laboratory within 10 min. The serum was separated by centrifugation at 4°C at 3000 rpm for 30 min and stored at −80°C until assayed for hormonal analysis and cholesterol esters (CE) profile. Follicle-stimulating hormone (FSH), luteinizing hormone (LH), thyroid stimulating hormones (TSH), porcine prolactin (PRL), anti-müllarian hormone (AMH), and testosterone were measured by enzyme-linked immunosorbent assay (Shanghai Enzyme-linked Biotechnology Co., Ltd.). The serum progesterone (P4) and estradiol (E2) concentrations were measured using a radioimmunoassay in the Beifang Institute of Biological Technology (Beijing, China). The serum leptin concentrations were measured using Porcine Leptin Elisa Kit (SEKP-0278, Solarbio life sciences, China).

Serum samples lipid extraction, separation of cholesterol esters and phospholipids via thin-layer chromatography (TLC), and determination of fatty acids via gas chromatography were conducted as described previously [20]. The fatty acid composition was analyzed with An Agilent 7890A gas chromatographer equipped with 15 m × 0.25 mm × 0.25 μm DB-WAX columns (Agilent). While, C15:0 was used as a standard for quantitation. All the data were first normalized with the phospholipids (PL) profile and then presented as mean ± SD.

### Luteal Cell Culture

Fresh ovaries of leptin and WT pigs were removed by laparotomy and immersed immediately in sterile phosphate buffer saline (PBS) with 5% penicillin-streptomycin solution and transported to laboratory. The luteal tissue were obtained from the ovaries and cut into small pieces using sterilized ophthalmic scissors, minced well and dispersed by pipetting in M199 (Sigma, M4530) medium containing collagenase (10 mg/mL) and the tissue was digested on a horizontal shaker in an incubator at 37°C for 30 min. The supernatants containing luteal cells were then decanted through nylon mesh to remove debris. The cells were washed twice by centrifugation and resuspended in 10 ml Dulbecco’s modified Eagle’s medium (DMEM) with 10% fetal bovine serum (FBS) and incubated at 37°C, 5% CO_2_ in a 96-well plate for 24 hours and then cells were collected and used for qPCR.

### Quantitative Polymerase Chain Reaction (qPCR)

Total RNA from ovarian tissue and luteal cells was isolated using the Trizol reagent (Transgen Up, China) according to the manufacture’s instruction. cDNA was synthesized from total RNA using a PrimeScript RT reagent Kit (TaKaRa, Japan) and was used as a template to perform qPCR in SYRB green-based qPCR instrument (CFX-96, Bio-Rad, USA). The reaction was performed in a 20 μL reaction mixtures comprising 10 μL of 2× SYBR (TaKaRa, Japan), 1 μL of cDNA, 1 μL of forward primer, 1 μL of reverse primer, and 7 μL of ddH2O. The reaction program as following: 95°C for 30 s, followed by 40 cycles of 95°C for 10 s and 62°C for 45 s. The relative expression levels of target genes were quantified by 2-ΔΔCt. The primers are listed in Table S2.

### Ovarian Histology and Immunohistochemistry

For histological examination, ovaries were fixed in 4% paraformaldehyde for 48∼72 h, processed by an automatic tissue processor (Yd-12p, Jinhua Yidi, medical appliance Co., Ltd, Jinhua, China) and embedded in a paraffin block (Yd-6D, Jinhua Yidi, medical appliance Co., Ltd, Jinhua, China). The paraffin blocks were cut into 5-um- thick sections using a Microm HM 325 microtome (Thermo Scientific, Waltham, MA, USA) and allowed to dry on glass slides overnight at 37°C. Thereafter, the tissue sections were deparaffinized in xylene and rehydrated through graded ethanol dilutions. Sections were stained with hematoxylin–eosin (H&E) (G1120; Solarbio, China) according to manufacturer’s instruction.

For immunohistochemistry (IHC), after dewaxing and hydration, sections were incubated in 3% H_2_O_2_ solution for 30 min, and washed with PBS for thrice (each time 3 min). After that, the sections were blocked in PBS containing 5 % BSA for 15 min at room temperature. Finally, the tissue sections were incubated with caspase-3 antibody (Table S3) at 4°C overnight. After washing with PBS for thrice, sections were incubated with specific secondary antibodies (R&D, USA) for 20 min. After washing thrice again, sections were stained with fresh DAB (KIT-9901, Elivision TM plus Polyer HRP IHC Kit, Fuzhou, China) solution in dark for 5 min and the washing with PBS thrice (each time 3 min). Hematoxylin counter-staining, and neutral gum sealing slides. Imaged using OLYMPUS BX53 fluorescence microscope and analyzed using software of accessories.

### Protein Extraction and Immunoblotting

The ovarian tissue from WT and leptin pigs was used to evaluate the expression of different protein levels (listed in Table S3) using western blotting. In brief, the ovarian tissues were lysed in RIPA lysis buffer (Bestbio, China) with protease inhibitors at 4 °C. After lysis, supernatants were obtained by centrifugation at 13000 × g for 15 min at 4 °C. Equal amounts of protein (100 μg) were run on SDS-PAGE gel, along with molecular weight marker. After electrophoresis, the proteins were transferred to PVDF membranes and reacted with primary antibodies against various antibodies and *β-actin* at 4 °C overnight. After incubation, the membranes were washed and incubated with specific secondary antibodies (R&D, USA). The membranes were then incubated with ECL (Easysee Western Blot Kit, China) and visualized with an Imaging System (Bio-Rad, Universal Hood II, USA).

### RNA-Seq Analysis and Quality Assessment

Total RNA was extracted using TRIzol Reagent (Invitrogen, CA, United States) and purified using an RNeasy Mini Kit (Qiagen, CA, United States). The quality of RNA was assessed using an Agilent 2100 Bioanalyzer (Agilent, Palo Alto, CA, United States). The rRNA-depleted RNA samples were further processed in accordance with the Illumina protocol (New England Biolabs, Massachusetts, United States). After cDNA synthesis, the samples were sequenced with an Illumina Novaseq6000 by Gene Denovo Biotechnology Co. (Guangzhou, China). The raw data were recorded. The overall quality of the RNA−seq data was evaluated by fastp (version 0.18.0). Clean reads were aligned to the Sus scrofa reference version 11.1 using HISAT2. 2.4 [21] with the default parameters.

### Screening and Clustering Analysis of Differentially Expressed Genes

Data preprocessing and follow-up analysis were performed DESeq2 software. The lists of DEGs between WT and leptin pigs cases were generated using the edgeR package (version 3.32.0) (Robinson et al., 2010). To normalize the raw data, the genes with the parameter of false discovery rate (FDR) below 0.05 and absolute fold change≥2 were considered differentially expressed genes between the leptin and WT groups. Hierarchical clustering analysis was performed based on the expression levels of all transcripts and significantly differentially expressed transcripts using the pheatmap R package based on Euclidean distance.

### Functional Enrichment Analysis

The Kyoto Encyclopedia of Genes and Genomes (KEGG) pathway enrichment analysis in each module and network was conducted using the Database for Annotation, Visualization and Integrated Discover (DAVID). DEGs and enriched pathways were mapped using KEGG pathway annotation with KOBAS3.02. The top 5 KEGG pathways were selected and ranked by the enrichment factor. Subsequently, the Venn diagram tool was used to help identify the common genes that were the focus of this work. Finally, gene set enrichment analysis (GSEA) using software GSEA and MSigDB [22] to identify whether a set of genes in specific GO terms.

### Single Nucleus Transcriptome Sequencing (Snrna-Seq) Libraries Construction

Cellular suspensions were loaded on a 10X Genomics GemCode Single-cell instrument that generates single-cell Gel Bead-In-EMlusion (GEMs). Libraries were generated and sequenced from the cDNAs with Chromium Next GEM Single Cell 3’ Reagent Kits v2.following the manufacturer’s instructions.

### Single-cell RNA-seq Data Processing

FASTQ files of 2 samples were processed with the use of Cell Ranger (v.3.1.0) count pipeline coupled with *Sus scrofa* reference version 11.1 to generate feature-barcode matrices. Seurat object list was then generated by Seurat package (v.4.0.4) [23], with R software (v.4.1.0) following these criteria: (1) min.cells = 5; (2) 200 < nFeature_RNA < 3600; (3) percent.mt <0.1. In other words, genes expressed in at least 5 cells and gene number detected in cells ranging from 800 to 10,000 were kept for further analysis, and low-quality cells were filtered if R 10% UMIs derived from the mitochondrial genome.

### Dimensionality Reduction and Clustering

To remove batch effects across samples, canonical correlation analysis (CCA) method was used for data integration [24]. In detail, we initially normalized the filtered gene expression data with Normalize Data function with parameter (normalization.method = “LogNormalize”, scale.factor = 10000) Then top 2,000 variable genes were identified using the ‘vst’ method with FindVariableFeatures function in each sample, respectively. At last, we used FindIntegrationAnchors with parameters (ie anchor.features = 2000, k.filter = 200, dims = 1:30) and IntegrateData functions with the top 30 dimensions among 2 samples. After filtering low quality cells with 200 < nFeature_RNA < 3600, percent.HB < 5 and nCount_RNA < 10000, we ran ScaleData function. To perform dimensionality reduction, the RunPCA function was conducted on linear-transformation scaled data with 2000 variable features and we performed UMAP with the top 30 dimensions. Finally, we clustered cells by using the FindNeighbors and FindClusters (resolution = 0.5) functions, which got 16 clusters.

### Cell-type Annotation and Cluster Marker Identification

Cell clusters were obtained, after dimensional reduction and projection of all cells into two-dimensional space by UMAP. The Seurat FindAllMarkers function was used to identify markers for each cell cluster with the default settings. Canonical markers of specific cell types were used for cluster annotation. Ovarian cell-type markers were selected from PanglaoDB database(https://panglaodb.se/), CellMarker database(http://biocc.hrbmu.edu.cn/CellMarker/) and literature review.

### DEG Identification and Functional Pathways Enrichment

The FindMarkers function in Seurat was used to identify differential expressed genes (DEGs) between two groups of cells with default parameters (logfc.threshold = 0.25, test.use = ‘‘wilcox,’’ min.pct = 0.1), and the enrich GO function in the clusterProfiler package (v.3.15.2) [25], was used to perform functional analysis with differential gene sets annotated with GO database (http://geneontology.org/). Enrichment pathways were obtained with parameters (pvalueCutoff = 0.05, pAdjustMethod = “BH”, OrgDb = “org.Ss.eg.db”).

### Cell Trajectory Construction by Monocle

The subpopulation of germ line or granulosa cell line was imported into Monocle (version 2.20.0) to dissecting cell differentiation fate, also termed “pseudotime analysis.” With the gene count matrix as input, the new dataset for Monocle object was created, and functions of “reduceDimension” and “orderCells” were carried out to generate the cell trajectory based on pseudotime. Particularly, the ordering genes were differentially expressed genes between clusters in each cell type calculated by “differentialGeneTest” function in Monocle. In addition, the root state (that is, a prebranch in the heatmap) was set and adjusted following consideration of the biological meanings of different cell branches.

### The Regulon Activity of Transcription Factors with SCENIC

The SCENIC algorithm was utilized to analyze the activity of transcription factors (TFs) and identify regulons (TFs and their target genes) in individual cells. The gene expression matrix, with genes in rows and cells in columns, was input into SCENIC (version 0.9.1) [26]. Co-expressed genes for each TF were constructed using GENIE3 software, followed by Spearman’s correlation analysis between the TFs and their potential targets. The “runSCENIC” procedure was then employed to generate the Gene Regulatory Networks (GRNs), also known as regulons. Regulon activity was analyzed using the AUCell (Area Under the Curve) software, applying a default threshold to categorize specific regulons as “0” (off) or “1” (on). t-SNE parameters were set to visualize the data with 50 principal components (PCs) and a perplexity of 50, determined through a consistency test across multiple perplexity values and number of PCs. Cell states were mapped using specific regulons, and the average binary regulon activity was calculated.

### Cell–cell Communication Analysis

Cell–cell communication was analyzed using iTALK package. The input data were Seurat object, and the ligand–receptor pairs were detected from top 50% highly expressed genes. The communication types mainly included growth factors, cytokines, and checkpoint. The network plot was visualized using LRPlot function.

### Statistical Analysis

Analyses were performed in the SAS statistical software (SAS ver. 9.4, SAS/STAT, SAS Institute Inc., Cary, NC). Data were expressed as least square mean ± standard error of mean unless otherwise stated. Continuous variables and measured over time, such as body weight (BW) and size including length, height, heart girth, chest depth, chest width and abdominal circumference, and hormones were analyzed by using the MIXED procedure of SAS (SAS/STAT) with estimations carried out by the method of restricted estimated maximum likelihood. All the mixed models included the repeated statement as day or stage of the estrous cycle and random effect of the animal identification nested within treatment for the proper error term. Continuous variable but measured on single point, such as serum leptin levels, pubertal onset, inter-estrus interval, back fat thickness, serum circulating cholesterol esters profile, relative expression of mRNA and protein and underweight infertile human serum profile were analyzed by PROC TTEST in SAS. Statistical significance was defined as \**P*<0.05, \*\**P*<0.01.

## RESULTS

### Screening of Leptin Pigs and Growth Performance

To evaluate the effect of leptin overexpression on female reproduction, leptin overexpressing transgenic sows were generated by breeding the wild-type (WT) sows with already generated leptin overexpressed male [18]. In brief, leptin overexpressed male was re-cloned and the cloned offspring (11) were mated with Duroc/Landrace/Yorkshire sow (DLY, 11) to obtain the F1 generation, and then the F1 generation was further crossed to obtain the F2 generation and so on (Fig. 1. A). The overexpression of leptin in offspring was confirmed by PCR and circulating serum levels of leptin (Fig. 1. B and C). The offspring-harboring transgenic vector (leptin gene with EGFP) were designated as leptin pigs, while, the others lacking transgenic vector were defined as WT group (Fig. 1. B). The leptin expression in leptin pigs was further elucidated by high levels of serum leptin (3045.9±857.9 vs 521.1±109.6 pg/mL, *P*<0.05; Fig. 1. C). To assess the growth characteristics of leptin overexpressing pig, we documented the growth parameters, including body weight, length, height, heart girth, chest width, chest depth and abdominal circumference (Fig. 1. D-J). The results showed that leptin pigs displayed slow growth rate as their body weight was significantly lower than WT pigs as at 180 days of age (30.0±2.4 vs 101.8±3.8 kg, *P*<0.05, Fig. 1. D). Nevertheless, the leptin pigs gradually gained weight but they remained underweight as compared to WT pigs (*P*<0.05). Similarly, the other growth parameters like length, height, heart girth, chest width, chest depth and abdominal circumference were also significantly lower in leptin pigs as compared to WT (*P*<0.05; Fig. 1. E-J). Additionally, the back fat thickness in leptin pigs was significantly lower than WT (5.6±6.4 vs 42.9±1.9 mm, *P*<0.05; Fig. 1. K). Collectively, these findings indicated that leptin pigs exhibited stunted growth with decreased back fat thickness.

**Fig. 1:**
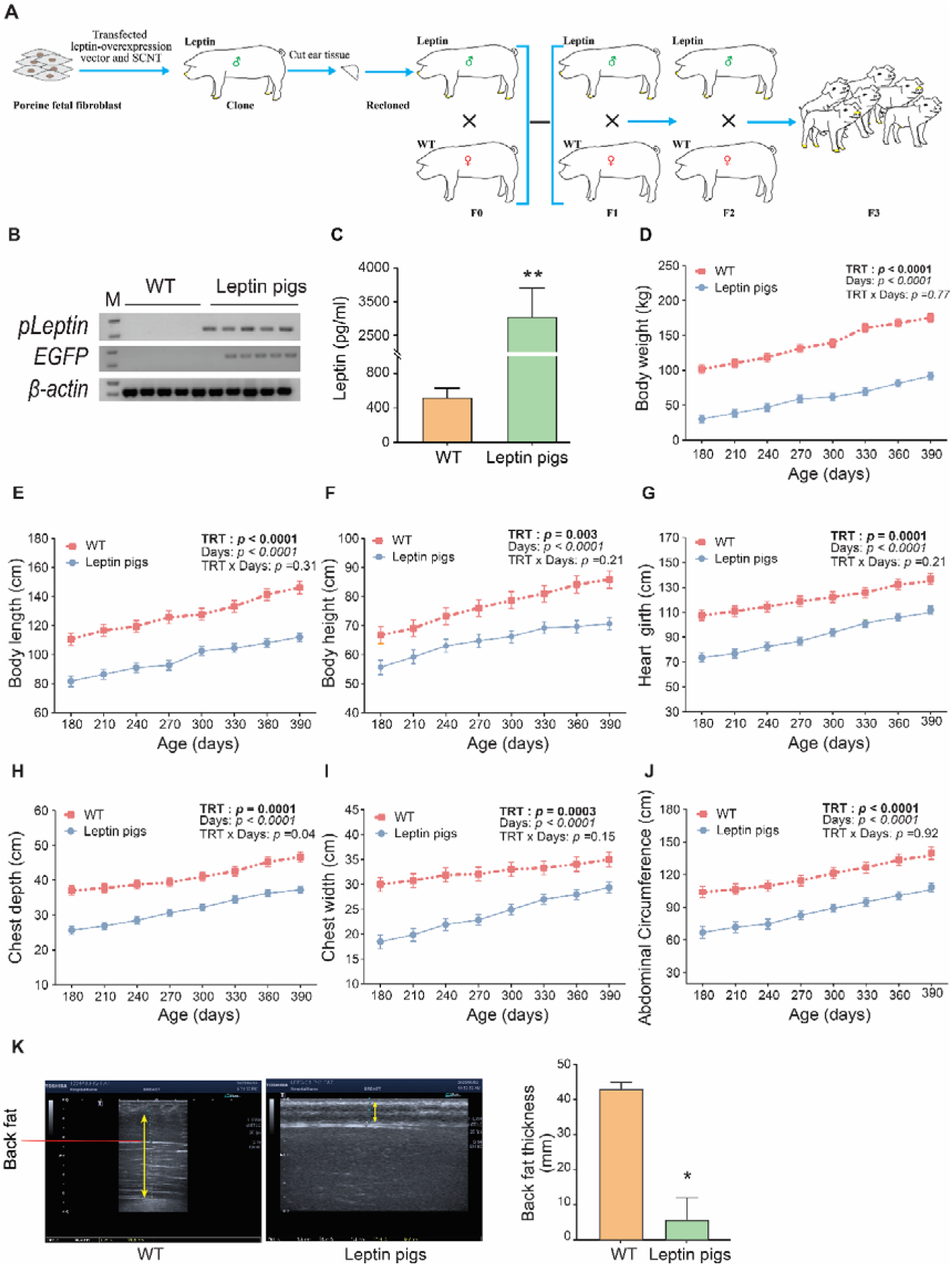
Screening of leptin pigs and body development: **(A)** Schematic diagram of leptin pigs’ generation and breeding **(B-C)** Screening of leptin pigs: **(B)** Leptin and EGFP expression was confirmed by 1% AGE. Lane M indicates DNA marker (DL2000). *β-actin* was used as an internal control **(C)** Serum leptin concentration in WT (n=4) and leptin pigs (n=4) **(D-J)** Changes in body weight and size comprising of length, height, heart girth, chest depth, chest width and abdominal circumference between WT (n=4) and leptin pigs (n=5) **(K)** Measurement of back fat thickness by ultrasonography in WT and leptin pigs, The back fat thickness in leptin pigs was significantly low (*p*<0.05), WT (n=3) and leptin pigs (n=3).

### Reproductive Performance of Leptin Pigs

Leptin has been involved in regulation of female reproduction and we evaluated the reproductive performance of leptin pigs comprising of pubertal onset, estrus expression pattern and breeding performance (Fig. 2). It was observed that the onset of estrus in leptin pigs (n=8) was at the age of 294d, 416d, 259d, 407d, 335d, 352d, 290d and 287d respectively. Collectively, the leptin pigs showed delayed onset of puberty as compared to WT pigs (330.0±54.3 vs 155±14.7 days, *P*<0.05; Fig. 2. A). Next, we evaluated the pattern of estrus expression in both leptin and WT pigs and observed that leptin pigs manifested irregular estrous expression characterized by both long and short cycles as sometimes they displayed estrus as short as after 9 days and sometimes as long as after 63 days (Fig. 2. B). The average length of estrous cycle in leptin pigs (n=8) was 35.8d, 40d, 28.9d, 33d, 23.4d, 23d, 24.5d and 25d respectively, while in WT pigs (n=5) it was 21.4d, 22.4d, 21.3d, 20.6d and 21d respectively. Collectively, the inter-estrous interval in leptin pigs was significantly higher than WT (29.2±6.0 vs 21.3±0.7 days, *P*<0.05; Fig. 2. C). Meanwhile, we started the breeding of leptin pigs (n=4) and it was observed that leptin pigs received more mating until they got pregnant. As two sows were bred for 3 times and at 3^rd^ breeding, they became pregnant (Fig. 2. D & E). While, the 3^rd^ leptin sow was bred for 6 times (Fig. 2. F), and 4^th^ sow was bred for 5 times (Fig. 2. G), but both heads remained non-pregnant (Fig. 2. F & G). Collectively, these data indicated that leptin pigs exhibited a greater number of mating (17 times) with a minimum of 3 times to a maximum of 6 times mating. Contrarily, WT pigs (n=10) were mated 10 times, and out of that, 2 times they failed to get pregnant, while 8 times they exhibited successful pregnancy. Whilst, the leptin pigs mated 17 times, out of which 15 times they remained non-pregnant while 2 times they became pregnant (Fig. 2. H). Thus, delayed onset of puberty and irregular estrous expression was followed by breeding insufficiency as a greater number of mating in leptin pigs were required until successful pregnancy.

**Fig. 2:**
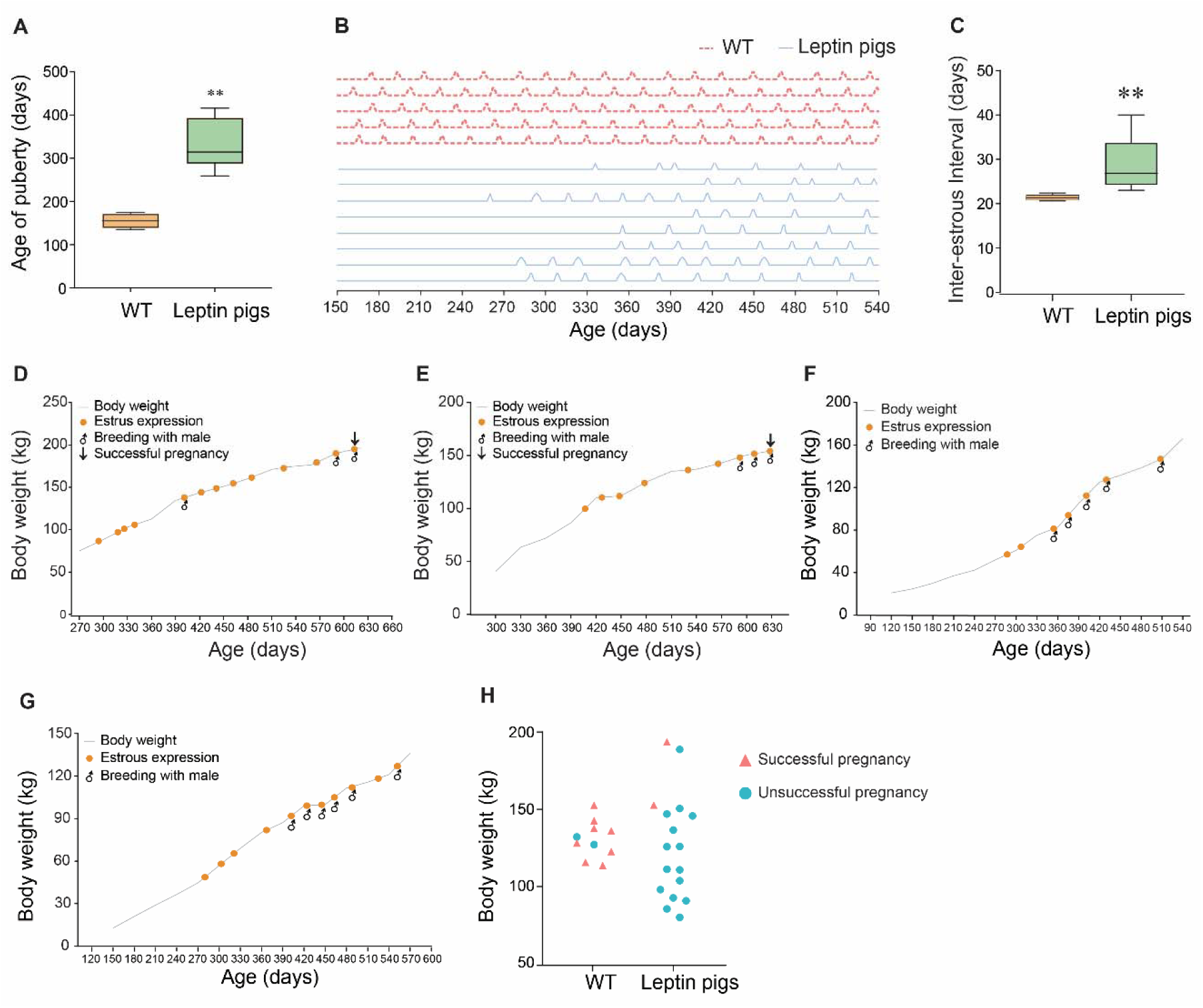
Puberty, estrus expression and breeding performance of leptin pigs. **(A)** Age of puberty in WT (n=5) and leptin pigs (n=8) **(B)** Pattern of estrus expression in WT (n=5) and leptin pigs (n=8). Each peak represents the estrous expression. **(C)**. Comparison of inter-estrous interval between WT (n=5) and leptin pigs (n=8). **(D)** Breeding performance of individual leptin pigs (n=4). Two heads (D,E) get pregnant at 3^rd^ time of breeding with male, while other 2 heads (E,F) remain non-pregnant even after 5-6 times breeding. **(H)** Comparison of breeding efficiency between leptin (n=10) and WT (n=4) pigs.

### Hormonal and Lipid Profiles of Leptin Pigs and Steroidogenesis

Gonadotropins and steroids regulate the reproductive axis and therefore, we evaluated the serum hormonal profile in leptin pigs at five different stages of the estrous cycle. Results showed that the concentrations of FSH, E2, TSH, and PRL in leptin pigs were higher (*P*<0.05) as compared with WT regardless of the estrous cycle stage (Fig. 3. A & B, E & F). Further, the concentrations of LH and P4 were also significantly higher in leptin pigs but interaction (*P*<0001) between treatment and the estrous stage was observed (Fig. 3. C & D) that indicated that the concentration of LH in WT pigs almost remained consistent across all stages of the estrous cycle, however, in leptin pigs it increased in early stages of the estrous cycle and then declined at the end of estrous cycle (d12 post met-estrus). Despite this, the concentrations of P4 were similar at earlier stages of the estrous cycle (proestrus, estrus, and met-estrus) in both leptin and WT pigs but it was significantly higher in leptin pigs than WT during d6 and d12 after met-estrus (*p*<0.05; Fig. 3. D). However, the concentrations of testosterone and AMH hormones remain unchanged in leptin pigs (*P*>0.05; Fig. 3. G & H). Collectively, these results indicated that the irregular estrous in leptin pigs followed an abnormal hormonal profile.

**Fig. 3:**
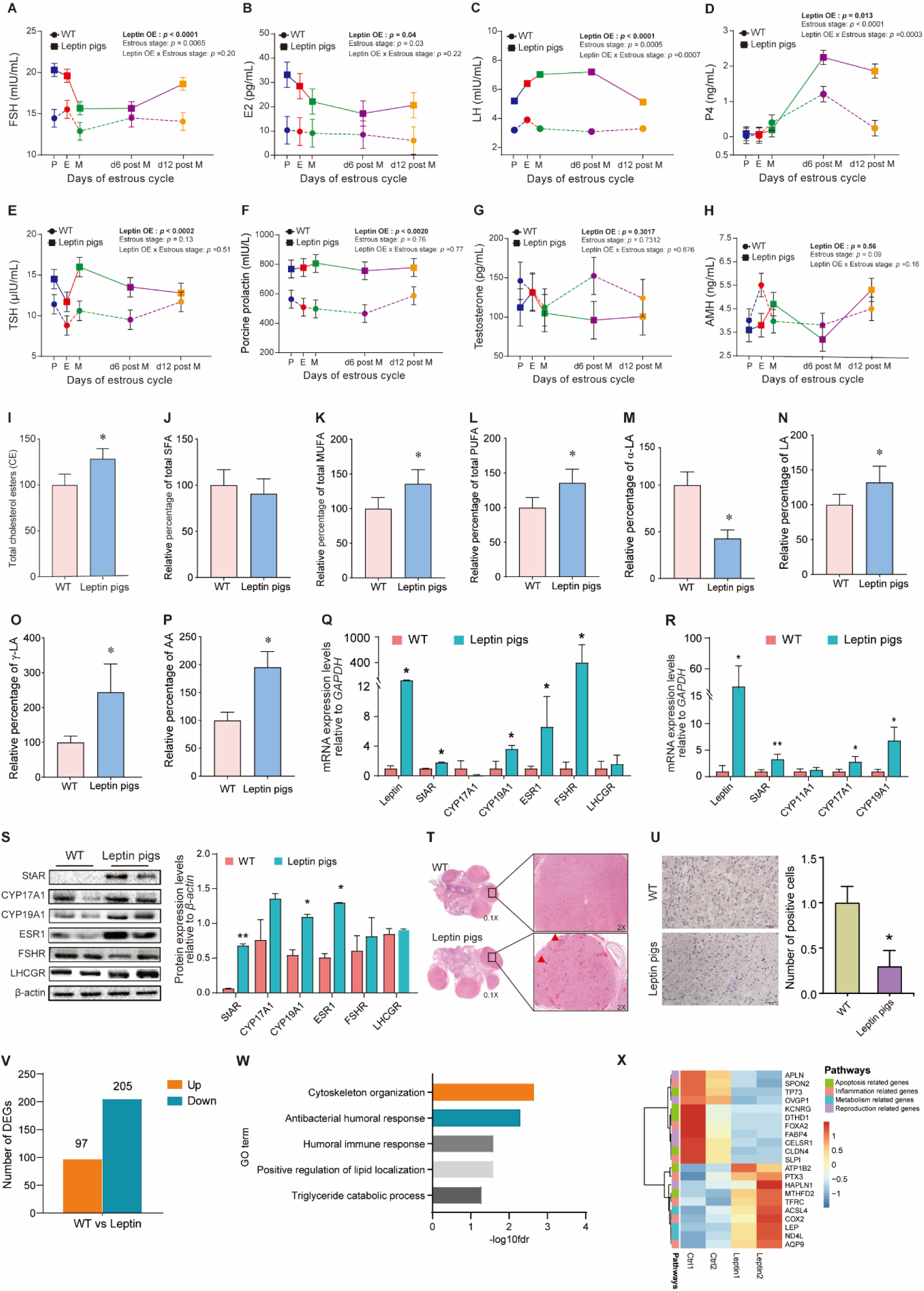
Serum hormonal changes in leptin pigs and steroidogenesis: Serum hormonal changes at five different stages of estrous cycle comprising of **(A)** follicle stimulating hormone (FSH) **(B)** estradiol (E2) **(C)** luteinizing hormone (LH) **(D)** progesterone (P4) **(E)** thyroid stimulating hormones (TSH) **(F)** porcine prolactin (PRL) **(G)** testosterone and **(H)** anti-müllerian hormone (AMH) levels in leptin (n=4) and WT (n=4) pigs. P: proestrus; E: estrus; M: metestrus; d6 post M: day 6 post metestrus and d6 post M: day 12 post metestrus. Leptin OE stands for leptin overexpression. **(I)** Serum levels of total cholesterol esters (CE) **(J)** Total saturated fatty acids (SFA) **(K)** Total monounsaturated fatty acids (MUFA), **(L)** Total polyunsaturated fatty acids (PUFA) in leptin and WT pigs. **(M)** Serum levels of α-linolenic acid (α-LA), **(N)** linolenic acid (LA), **(O)** γ-linolenic acids (γ-LA) and **(P)** arachidonic acids (AA) in WT (n=3) and leptin pigs (n=3). **(Q)** mRNA expression of steroidogenic markers at ovary of WT (n=2) and leptin pigs (n=2). *GAPDH* was used as an internal control. **(R)** mRNA expression of steroidogenic markers at luteal cells. *GAPDH* was used as an internal control. **(S)** Protein abundance of steroidogenic markers at ovary of WT (n=2) and leptin pigs (n=2). *β-actin* was used as an internal control. **(T)** Ovarian histology of WT and leptin pigs. **(U)** The expression of apoptotic markers at ovary of WT and leptin pigs. The number of cas-3 positive cells were calculated after IHC. \**P<0.05*, ** *P<0.01*. **(V)** The number of differentially expressed genes in leptin and WT group. **(W)** Gene Ontology and Kyoto Encyclopedia of Genes and Genomes (KEGG) analysis performed by Metascape. (**X)** Heatmap of genes enriched in different pathways.

To clarify the essence of body fat composition on reproductive performance, we further assessed the blood CE profile in WT and leptin pigs. It was observed that serum total CE levels in leptin pigs were significantly higher than in WT pigs (125.6±17.11 vs 100.00±12.5, *P*<0.05; Fig. 3. I). Levels of total saturated fatty acids (SFA) in leptin pigs remain unchanged (90.8±16 vs 100.0±16.8, *P*>0.05; Fig. 3. J), while the levels of monounsaturated fatty acids (MUFA) and polyunsaturated fatty acids (PUFA) were increased in leptin pigs as compared with WT (136.0±20.1 vs 100.00±16.0, *P*<0.05; Fig. 3. K), and (136.0±19.5 vs 100.0±14.6, *P*<0.05; Fig. 3. L), respectively. Specifically, α-linolinic acid (α-LA) levels in leptin pigs were significantly decreased (42.9±8.9 vs 100.0±14.2, *P*<0.05; Fig. 3. M). While, leptin pigs also exhibited *(P* <0.05) high levels of linolenic acid (132.1±23.2 vs 100.0±15.0), γ-linolenic acid (244.3±80.8 vs 100.0±17.8), and arachidonic acid (195.5±28.0 vs 100.0±14.7) than WT pigs (Fig. 3. N-P). Collectively, the levels of CE (precursors of steroids) were increased in leptin pigs.

We evaluated the mRNA and protein expression patterns of steroidogenic markers. Results indicated that mRNA expression of leptin, steroidogenic acute regulatory protein (StAR), cytochrome P450 19A1 (CYP19A1), estrogen receptor 1 (ESR1), and follicle-stimulating hormone receptor (FSHR) were higher (*P<0.05*) in leptin pig ovaries, while the mRNA expression of cytochrome P450 17A1 (CYP17A1) and luteinizing hormone/choriogonadotropin receptor (LHCGR) was comparable between leptin and WT pigs (*P*>0.05, Fig. 3. Q). Likewise, the mRNA expression of leptin, *StAR*, *CYP17A1,* and *CYP19A1* was significantly higher (*P<0.05*) in leptin luteal cells, while *CYP11A1* was comparable (*P*>0.05) between leptin and WT luteal cells (Fig. 3. R). Further, western blot analysis indicated that the protein abundance of StAR, CYP19A1 and ESR1 was significantly higher in leptin pigs’ ovaries (*P*<0.05, Fig. 3. S), while CYP17A1, FSHR and LHCGR abundance was comparable between leptin and WT pigs (*P*>0.05, Fig. 3. S).

Although histological analysis revealed there was no significant difference in the number of early follicles and sinus follicles in leptin transgenic pigs relative to WT. However, leptin pigs exhibited more severe vascular inflammatory infiltration of the ovarian corpus luteum relative to the corpus luteum of WT pigs (Fig. 3. T). Further, the IHC analysis of ovaries revealed that the number of Cas-3 positive cells was significantly lower in leptin pig ovaries than in WT ovaries (P<0.05, Fig. 3. U). Collectively, these results indicated that the enhanced expression of StAR and CYP19A1 at tissue and cell levels was followed by inflammatory changes and compromised apoptosis in leptin pig ovaries.

### Transcriptome Profile and GSEA of Pig Ovaries

Our RNA sequencing analysis uncovered significant differences in gene expression between the leptin-overexpressing (leptin) group and WT group, with a total of 312 differentially expressed genes. Among these 97 were upregulated, while 205 were downregulated in leptin group (Fig. 3. V, Fig. S1. A). To gain insight into the functional implications of these differentially expressed genes, we conducted functional enrichment analysis, which revealed several significantly enriched processes. These included cytoskeleton organization, antibacterial humoral response, humoral immune response, positive regulation of lipid localization, and others (Fig. 3. W). Furthermore, gene set enrichment analysis (GSEA) revealed negative regulation of interleukin-2 production pathways was downregulated in leptin pigs (Fig. S1. B). To visualize the differentially expressed genes, a heatmap was generated, and some genes crucial for apoptosis (TP73, CLDN4, DTHD1, and KCNRG) and reproduction (APLN, OVGP1, FABP4, and CELSR1) were downregulated in leptin pigs. Whilst, the genes involved in metabolic changes (ACSL4, LEP, and ND4L) and inflammatory changes (PTX-3, TFRC, COX-2 and AQP9) were upregulated in leptin pigs (Fig. 3. X). Collectively, these data revealed that the leptin pigs’ ovaries exhibited inflammatory changes and compromised apoptosis and reproduction, accompanied by notable metabolic alterations. These insights contribute to our understanding of the molecular mechanisms underlying the effects of leptin overexpression on ovarian function.

### Single-nucleus Transcriptome Profiling of Pig Ovaries Identified Different Ovarian Cell Types and Gene Expression Signatures

To gain a deeper understanding of the impact of high leptin levels on individual ovarian cells, we characterized the distinct cell lineage and gene expression dynamics in both WT and leptin-overexpressed pigs. This was achieved by performing single-nucleus RNA-seq (snRNA-seq) on the ovarian tissue of both wild-type *sus scrofa* and our transgenic model, which were then analyzed using bioinformatics tools (Fig. S2. A). After stringent cell filtration, the high-quality transcriptomes of 17108 single cells (8686 in the WT group and 8422 in the leptin group; Fig. S2. B) were retained for subsequent analyses. We performed unsupervised clustering and grouped the total ovarian cells into 11 clusters as visualized in the Uniform Manifold Approximation and Projection (UMAP) plot (Fig. 4. A). By using established marker genes of ovarian cells, we identified 11 clusters as large granulosa cells (LGC), theca-interstitial cells (TIC), luteal cells (LC), endothelial cells (EC), small granulosa cells (SGC), macrophage, lymphocytes (1 and 2), perivascular cells (PCs), and epithelial cells (EC). We found that both the leptin and WT group contained all kinds of cell types, while the cell fraction was different between groups (Fig. 4. B, Fig. S2. C).

**Fig. 4.**
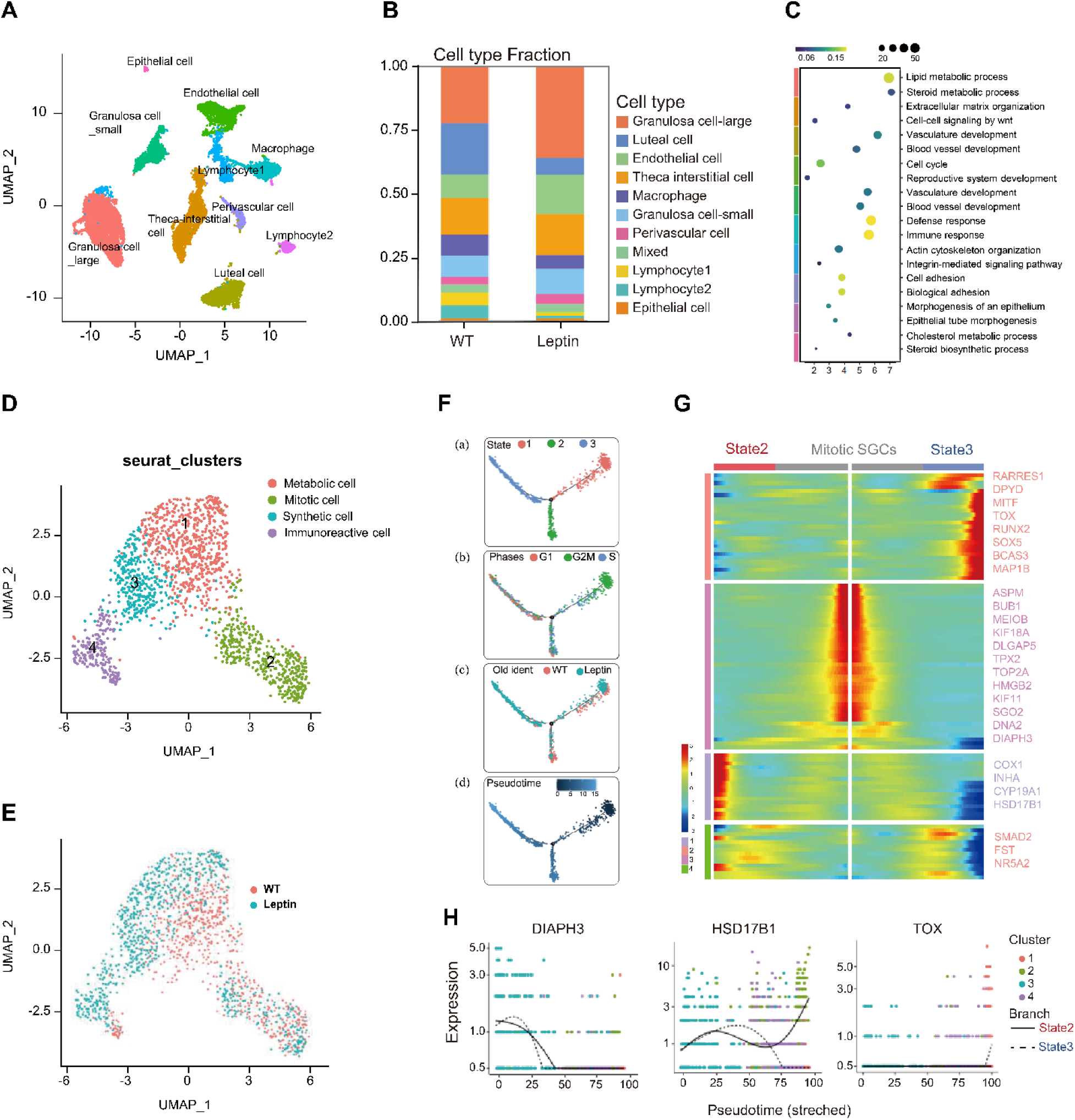
Single-Nucleus Transcriptome Profiling of Pig Ovaries Identified Different Ovarian Cell Types and Gene Expression Signatures. **(A)** UMAP plot of the merged dataset of leptin and WT pig ovaries showing 10 ovarian cell types. **(B)** Cell type fractions in WT and leptin group. **(C)** Representative GO terms of upregulated DEGs between ovarian cell types. Color in left is corresponding to Fig. 4.A to represent cell types. GeneRatio is showed by color from blue to yellow, Gene Count is indicated by size of circle. **(D)** UMAP plot in four subtypes of small granulosa cells **(E)**. UMAP plot shows WT and leptin cell distribution in SGC’s subtypes**. (F)** Single-cell trajectories of the 3 SGC states through the pseudotime (a). Single-cell trajectories along with cell cycle phase (b), WT and Leptin group distribution (c) and pseudotime (d). **(G)** Heatmap of two kinetic trends for state 2 and state 3. Middle line is the start of pseudotime.

All clusters were identified according to the specific expression of markers (Fig. S2. E). Clusters corresponding to granulosa cells (GCs) were identified based on the expression of granulosa cell markers, such as *CYP19A1* and *FST*. LGCs and SGCs were annotated according to their average RNA levels and INHA (an antagonist of granulosa cell proliferation) expression patterns. *NR5A2*, *RNF150* and *NLGN1* were specifically expressed at LGCs, while *INHA, MYH11* and SERPINE2 were specifically expressed at SGCs. Theca-interstitial cell markers *COL3A1* were specifically expressed in one cluster. Interestingly, some of cells in this cluster expressed high levels of the theca marker *PDGFRA*, implying that this cluster contained theca cells that are associated with the development and maturation of follicles [27] and *CDH11* and *SLIT2* were specifically distributed at TIC (Fig. S2. E). We next analyzed biological functions for every cell cluster by performing GO analysis of DEGs across all the clusters, revealing unique characteristics of these ovarian cells (Fig. 4. D). For example, GO terms of LGCs were involved in “lipid metabolic process” and “steroid metabolic process”. GO terms including “extracellular matrix organization” and “cell-cell signaling by wnt” were enriched in TIC, suggesting that TICs provide structural support and mediate signaling between ovarian cells. And GO terms including “blood vessel development” and “immune response” were enriched in ECs and macrophages, respectively. Collectively, we verified 10 different ovarian cell types in pig ovaries and depicted gene expression signatures for each cell type.

### Developmental Patterns of Small Granulosa Cells Indicate Inflammatory State in Leptin Pig

Granulosa cells play an important role in ovarian follicle development [28]. To investigate gene expression dynamics of SGCs, we performed unsupervised clustering and identified four subtypes of SGCs (Fig. 4. E). Notably, cluster 1 and 2 exhibit obvious overlap between WT and leptin groups, while cluster 3 and 4 were mainly consisted with cells from leptin group (Fig. 4. E). To further identify the clusters, we detected DEGs in comparisons between four SGC subpopulations and the highly variable expressed genes were visualized in UMAP plots (Fig. S3. A). For example, mitosis-associated genes, such as *DIAPH3*, *HMMR*, *SGO2*, and *CENPE* were mainly expressed in cluster 2 (green), while inflammation-related genes like *SOX5*, *TOX*, *AKAP12,* and *SOD* exhibited peak expression in cluster 4 (purple). Subsequently, GO analysis reveals biological variance between SGC subtypes (Fig. S3. B). For instance, GO terms including “androgen metabolic process” and “lipoprotein biosynthetic process” were enriched in cluster 1, genes involving “hormone biosynthetic process” and “steroid biosynthetic process” were enriched in cluster 3, GO terms including “mitotic nuclear division” and “regulation of cell cycle” were enriched in cluster 2, GO terms including “positive regulation of interleukin-2 production” and “cAMP catabolic process” were enriched in cluster 4. Therefore, we referred SGCs in subtypes as Metabolic cells, Mitotic cells, Synthetic cells, and Immunoreactive cells, respectively. In addition, the dominant population in Synthetic cells and Immunoreactive cells were mainly cells from the leptin group, indicating SGCs in leptin-overexpressed ovaries have distinct gene signatures regarding lipid metabolism and inflammation (Fig. 4. E).

To explore how SGC cell transition from one state to another, we constructed cell developmental trajectories by Monocle2. Cell pseudotime trajectories predicted 3 granulosa cell states and 1 branch point (Fig. 4. F). State1 was mainly consisted with cells from G2M phase, we assigned this state to Mitotic SGCs. In this stage leptin and WT cells were evenly distributed, showing similar transcriptional dynamics between two samples during early mitotic phase of SGCs. For further characterization of the small granulosa cell lineage, top 100 differentially expressed genes of the 3 states were divided in 4 clusters according to their distinct expression patterns, and a heatmap was generated (Fig. 4. G). Notably, state3 highly expressed SOX5, BCAS3, MITF and TOX that is essential for cell migration and immune response, showing state 3 is corresponding to immunoreactive cell. As state 3 is mainly consist of leptin cells, we confirmed the inflammation state in leptin pig occurs at later cell differentiation stage. Additionally, state2 exhibited sharply increased expression of metabolic genes, such as *COX1*, *INHA*, *CYP19A1* and *HSD17β1*, while mitotic SGCs had higher expression of *MEIOB*, *HMGB2* and *SGO2*. The gene plot (Fig. 4 H) showed two kinetic trends for state 2 or state 3, from branch point to the end of trajectories. *DIAPH3*, a cell mitotic gene, downregulated as cells went along the trajectory in both states, indicating loss of mitotic potential during developing process. We also showed *TOX*, an immune cell differentiation gene, which is upregulated during the developmental process in state 3. *HSD17β1* is a hormone metabolic enzyme; it is upregulated in state 2, indicating gain of metabolic function during normal development.

In order to describe the pre-transcriptomic profile of SGC subtypes, we further investigated regulons activity of transcriptional factors in SGCs using SCENIC (Fig. S3). Based on 125 regulons and 6048 filtered genes with default filter parameters, the cells were clustered by 4 subcluster identities of SGCs in t-distributed stochastic neighbor embedding (t-SNE) method calculated by SCENIC. The regulon activity was binarized and matched with cell identities and some representative regulons and their motifs were listed. As shown, *SOX5*, *SOX6*, *STAT5A* and *RUNX2* were active mainly in the Immunoreactive cells, while *IVD*, *MYBL1*, *E2F8* and *RAD21* were mostly active in Mitotic cells, indicating higher cell proliferation and signaling activities in these sub-clusters. Moreover, several TFs, such as *TBL1XR1* and *TCF4*, were significantly represented in all four sub-clusters, which supports the role of these TFs as likely candidates for sustaining SGC-specific transcription programs during all stages.

### Differential Expressed Genes Reveal Altered Hormone Metabolic Process in Leptin Pig

To further dissect the mechanism for high leptin-induced infertility, we compared all cells of the leptin group and WT group and chose the top 400 differential expressed genes to perform GO analysis. Our results revealed that the overall differential expressed genes were enriched in the lipid metabolic process and steroid metabolic process (Fig. 5. A), which are important for hormone homeostasis. This suggests that the leptin group may experience changes in hormone metabolism compared to the WT group. The GO term analysis revealed that the DEGs in leptin cells were mainly involved in steroid biosynthesis and other hormone synthesis-related pathways like cortisol, aldosterone and thyroid hormone synthesis, and metabolism-related pathways. While the DEGs in WT cells were enriched in tissue development and biological adhesion and cell differentiation pathways (Fig. S3. D). Collectively, these findings indicated that the DEGs in leptin cells exhibited mainly hormone biosynthesis and lipid metabolism-related pathways.

**Fig. 5:**
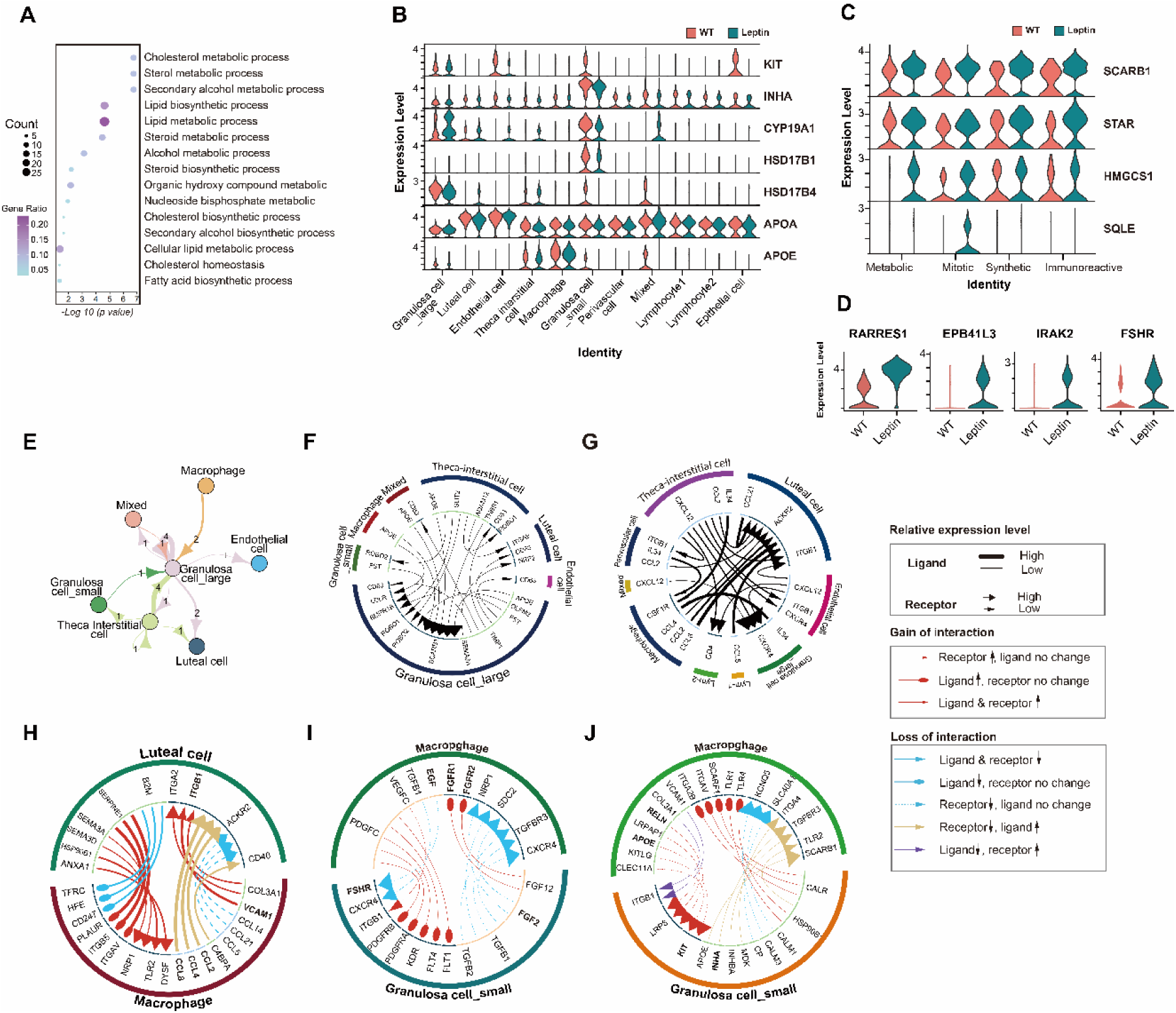
Differential expressed genes revealed altered metabolic process in leptin pigs. **(A)** Representative GO terms of DEGs between leptin and WT cells. **(B)** Vln plots of the expression level of representative differential expressed genes involved in reproductive and metabolic process across all cell population in in leptin and WT pig ovaries. **(C-D)** Vln plots of the expression level of genes involved in cholesterol biosynthesis in leptin and WT cells in four clusters of SGCs. **(E)** NetView plot showing interactions between different cell types. The width of edges represents the strength of the communication. Labels on the edges show exactly how many interactions exist between two types of cells. **(F-J)** Ligand receptor interaction between different cell populations. All cell types with all ligand-receptor categories **(F-G)**. Interaction of luteal cell and macrophage by top 20 differential expressed cytokines between leptin and WT groups **(H)**. Interaction of macrophage and SGCs through top 20 differential expressed growth factors **(I)** and other factors **(J)**. SCENIC results on SGCs. Cluster labels correspond to 4 subtypes in Fig4.D and cell state; TFs confirmed by literature (Regulons) and their corresponding enriched DNA-binding motifs are shown (Motifs).

To further evaluate the metabolic heterogeneity of small granulosa cells, we analyzed DEGs of SGCs between WT and leptin groups. GO terms including “cholesterol metabolic process” and “steroid metabolic process” were enriched in upregulated genes of the leptin group (Fig. 5. A). Interestingly, genes related to cell proliferation and metabolism such as *KIT*, *INHA*, *CYP19A1*, *HSD17B1*, and *HSD17B4* showed similar expression patterns across cell types. These were mainly expressed in granulosa cells and downregulated in the leptin group, suggesting granulosa cells are the main site regulating ovary development and function impairment in leptin pigs. While lipoprotein APOA was expressed in all cell types and slightly deceased in luteal cells and TICs, and APOE was downregulated in leptin macrophages (Fig. 5. B). Additionally, pro-apoptosis genes like RARRES1, EPB41L3, IRAK2, and reproductive hormone receptor FSHR were upregulated in the leptin group (Fig. 5. D). Further, genes participating in the cholesterol biosynthesis pathway, such as *SCARB1*, *STAR*, *HMGCS1*, and *SQLE* have a higher level in the leptin group (Fig. 5. C), indicating increased cholesterol biosynthesis. Strikingly, these genes were equally expressed at all four states of SGC except SQLE which was specifically exhibited in the mitotic state, suggesting a varying metabolic pattern during the differentiation of SGCs in the leptin group (Fig. 5. C). The results were in line with the phenotype observed in leptin pigs, suggesting disrupted hormone homeostasis and providing insights into the mechanisms underlying infertility in leptin pigs, particularly concerning cholesterol biosynthesis, granulosa cell function, and apoptosis regulation.

### Cell-cell Communication Indicates Inflammation of Corpus Luteum and Follicle Growth Arrest

To further explore the crosstalk among different clusters of pig ovaries, we established the cell-cell communication network of all cell clusters with the iTALK package (Fig. 5. E, F). Remarkably, the majority of ligand-receptor pairs were identified between large granulosa cells (LGCs) and theca-interstitial cells (TICs), indicating that the LGC and TIC complexes serve as the main hub for cell-cell communication. These cells were also found to exchange signals with other cell types such as macrophages and endothelial cells. Moreover, LGCs and TICs had crucial roles in sustaining steroid hormone metabolic homeostasis by APOE-SCARB1 ligand-receptor pairs and regulating follicle development via TIMP1-CD63, FST-BMPR1B, and SLIT2-ROBO1 ligand-receptor pairs. Additionally, when it comes to cytokines, nearly all the receptors were derived from luteal cells, and the associated ligands were enriched in immune cells including macrophages, lymphocyte cluster 1 (lym1), and lymphocyte cluster 2 (lym2), which suggests the important roles of luteal cells in ovarian immune response (Fig. 5. G).

Subsequently, we performed a comparative analysis of important cell-cell interactions such as those between macrophage and luteal cells, macrophage, and SGCs (Fig. 5. H-J). The analysis of cytokine communication between macrophages and luteal cells revealed upregulation of ligands such as CCL2, CCL4, and CCL8, along with receptors NRP1, TLR2, and DYSF in leptin-treated macrophages (Fig. 5. H). This finding suggests that leptin may mediate inflammation in the corpus luteum by activating macrophages. Additionally, the analysis of growth factor family interactions between macrophages and luteal cells showed upregulation of follicle-stimulating hormone receptor (FSHR) in leptin-treated SGCs, indicating a potential reduction in the FSH sensitivity of granulosa cells (Fig. 5. I). Furthermore, growth-associated factors including KIT, RELN, and INHA were downregulated in SGCs, implying inhibition of follicle growth (Fig.5. J). In a nutshell, the cell-cell communication analysis revealed that inflammation of the corpus luteum and follicle growth arrest may be attributed to altered interactions between macrophages and luteal cells, as well as changes in growth factor signaling within SGCs. These findings shed light on the complex interplay between different cell types in the ovary and provide insights into the potential mechanisms underlying the reproductive impairments observed in leptin-overexpressing pigs.

## Discussion

Leptin overexpressing transgenic female pigs had reduced body weight, growth, and fat depots along with delayed puberty, irregular estrous cycles, and greater number of matings. This was linked to metabolic imbalances, increased immune response, and altered ovarian functions. This study provided a theoretical basis for the complex mechanisms underlying leptin, and reproduction by employing leptin-overexpressed female pigs.

The constitutional delay in growth has a direct relation to leptin concentrations [29]. The leptin pigs exhibited delayed body weight and growth that coincides with the previous findings in mouse models overexpressing leptin [30] and is attributed to the anorexigenic effect of leptin, as chronic administration of leptin has been shown to reduce body weight and growth [10]. Previously, we observed that high leptin in pigs suppressed insulin and impacted glucose homeostasis. Therefore, this could be the possible mechanism behind the sluggish growth in leptin-overexpressed pigs [12]. Moreover, the leptin pigs also exhibited low back fat deposits, likely due to homeostasis imbalance in lipolysis and lipogenesis as evidenced by our previous study [18]. Therefore, the sluggish growth in these leptin pigs could be due to the anorexigenic and adipolytic activity of leptin.

Puberty is a fascinating developmental transition that gates the attainment of reproductive capacity and culminates in the somatic and sexual maturation of the organism [31]. Shreds of evidence from animal models and human studies suggested that leptin acts as a permissive factor for the onset of puberty and maintenance of reproductive cyclicity [11]. However, in our study, leptin overexpressing pigs displayed a late onset of puberty followed by irregular estrus comprising of short and long estrus and breeding insufficiency characterized by more mating. Contrarily, transgenic mice overexpressing leptin manifested accelerated puberty and maintained fertility at earlier ages, despite being lean like the leptin pigs [7]. This discrepancy may be attributed to species differences and threshold levels of leptin as several reports indicated that physiological doses of leptin influencing anorexigenic effect were not equally enough to boost reproductive functions as both functions of leptin are mediated via different hypothalamic pathways [11, 32, 33]. Therefore, it is also plausible to speculate that leptin pigs may not attain enough leptin to boost reproduction and hence exhibit reproductive impairment, however, such mechanisms need further exploration.

The initiation and maintenance of reproductive functions normally depend upon gonadotropins that in turn cause the production of steroids. However, endocrine hormonal imbalances due to inappropriate secretions of gonadotropins lead to ovulatory dysfunction that results in reproductive impairment [34]. In this study, we observed reproductive disorders in leptin pigs, characterized by elevated basal levels of gonadotropins (FSH & LH) and increased steroid production. This could be attributed to high leptin, as it is well-accepted that leptin also augments the secretions of gonadotropins [35]. According to the two-cell–two-gonadotropin theory, LH stimulates thecal-interstitial cells to produce androgens, and FSH stimulates granulosa cells to produce estrogens from androgens [36]. Our study showed both FSH and FSHR, and its target granulosa cell fraction was elevated in leptin pigs, which aligns with the elevated E2 level. We also observed elevated TICs and endothelial cells in leptin pig ovaries, corresponding to higher P4 and vascularization. This indicates the ovarian axis was overactivated in leptin pigs. It has been reported that elevated LH levels impair downstream ovarian folliculogenesis as the ‘FSH threshold’ required for follicle maturation is frequently not reached, causing follicles to arrest in a preovulatory stage, giving rise to cystic ovaries which results in menstrual irregularities [37]. This follicle arrest was also evident from the cell-cell communication network as KIT and INHA were downregulated in the SGC of the leptin group (Fig. 5). Enhanced production of KIT ligand has been shown to stimulate the oocyte growth through PI3K signaling pathways, while low expressions are insufficient to activate KIT on oocyte and thus maintained the oocyte in quiescent stage [38]. Further, a reduced luteal cell population, as well as elevated granulosa cells in leptin ovaries, pretends to impair granulosa cells’ transformation to luteal cells, which might impair ovulation. Furthermore, apoptosis is one of the necessary pathways for luteolysis, while downregulated apoptosis pathways in leptin pigs’ ovaries which might be due to reduced luteal cell count, also seem to impede the luteolysis which was also evident from high levels of P4 at the end of the estrous cycle. Thus, it is intriguingly speculated that anovulation and incomplete luteolysis may lead to irregular estrous in leptin pigs.

Steroids are synthesized from cholesterol esters (CE), highlighting the importance of cholesterol homeostasis for the maintenance of fertility. Disturbance in serum cholesterol can affect steroidogenesis and reproductive functions. In this study, leptin pigs exhibited high serum levels of CE which was also evident from snRNA sequencing that genes involved in cholesterol synthesis i.e. HMGCS1 and SQLE were upregulated in leptin ovaries. These high levels of CE along with the upregulation of genes mediating the transport of cholesterol from the outer mitochondrial membrane to the inner mitochondrial membrane i.e STAR and SCARB1 (Fig. 5C) pretend to increase steroidogenesis (P4) in leptin pigs. Further, α-LA has shown an inhibitory effect on cholesterol synthesis [39, 40], and arachidonic acid (AA) and its metabolites have long been implicated in steroidogenesis through direct effects on the steroidogenic machinery (e.g., acute steroid regulator [StAR] [41]. Thus, the low levels of α-LA, and high levels of AA along with enhanced expression of the regulatory protein (StAR) in leptin pigs indicate enhanced steroidogenesis which might be the cause of the estrous irregularity.

Macrophages play crucial and diverse roles in various intra-ovarian events, encompassing folliculogenesis and the formation and regression of the corpus luteum [42]. Our study reveals macrophages communicated with different ovarian cell types, including granulosa cells, theca-interstitial cells, and luteal cells, which are mediated by cytokine ligand-receptor interactions. On the one hand, the development of follicles relies on the presence of an appropriate cytokine and growth factor environment, and macrophages serve as a significant source of these factors. In leptin pigs, we observed that the SGC and macrophage interaction is biased in both the cytokine and growth factor categories. These findings coincide with the previous findings of PCOS patients in which GC expressed elevated transcripts encoding cytokines, chemokines, and immune cell markers [43]. On the other hand, macrophages play a pivotal role in various aspects of corpus luteum function. First, they contribute to angiogenesis within the corpus luteum by secreting important factors, our study showed increased VCAM1 and ITGB1 in macrophage-luteal cell communication of leptin pigs, indicating well-established vasculature in leptin pigs. This is also evidenced by the presence of rich vessels in the H&E section of leptin pig ovaries. Notably, the number of macrophages within the corpus luteum reaches its peak during regression, indicating their involvement in the process of luteolysis. Our leptin pig ovaries exhibited fewer macrophages than WT pigs, indicating that fewer luteal cells need to be eliminated. Therefore, we hypothesized that alterations in the macrophage-mediated regulation of folliculogenesis and the corpus luteum function contribute to the pathogenesis of ovarian disorders.

## Conclusions

In this study, we found that leptin overexpression in pigs adversely affects reproductive performance, causing stunted growth, delayed puberty, irregular estrous cycles, and reduced breeding efficiency. This is linked to metabolic imbalances, increased immune response, and altered ovarian functions. The study provides insights into the complex mechanisms underlying ovarian function and suggests a potential role for leptin in ovarian pathophysiology. These findings provided some theoretical basis for the reproductive role of leptin.

## List of Abbreviations

CYP19A1: Cytochrome P450 19A1
E2: Estradiol
ESR1: Estrogen receptor 1
FSH: Follicle-stimulating hormone
FSHR: Follicle-stimulating hormone receptor
GCs: Granulosa cells
GSEA: Gene set enrichment analysis
LC: Luteal cells
LGC: Large granulosa cells
LH: Luteinizing hormone
LHCGR: Luteinizing hormone/choriogonadotropin receptor
P4: Progesterone
PUFA: Poly unsaturated fatty acids
SFA: Saturated fatty acids
SGC: Small granulosa cells
snRNA-seq: Single-nucleus RNA sequencing
StAR: Steroidogenic acute regulatory protein
TIC: Theca-interstitial cells
WT: Wild type
α-LA: α-linolinic acid

## Declarations

### Ethics approval and consent to participate

Animals used in this study were regularly maintained in the Laboratory Animal Centre of Yunnan Agricultural University. All animal experiments were performed with the approval of the Animal Care and Use Committee of Yunnan Agricultural University.

### Consent for publication

Not applicable.

### Availability of data and materials

The dataset supporting the conclusions of this article is included within the article and its additional file. The raw data for snRNA sequencing are submitted as GSE237153.

### Authors’ Contributions

HJW conceived the project. HJW, HYZ, and RZ designed the research experiment. MAJ, YC, DJ, JG, HZ, TW, HL, WC, DZ, WX, SP and YC conducted experiments. HYZ, KX, DJ, MAJ, YQ and BJ acquired data. MAJ, YC, and DJ analyzed data. HJW, HYZ, MAJ, YC, KX and RZ wrote the manuscript.

## Acknowledgements

We thank the Department of Science and Technology of Yunnan Province for their financial support.

## Supplementary materials

**Fig. S1:**
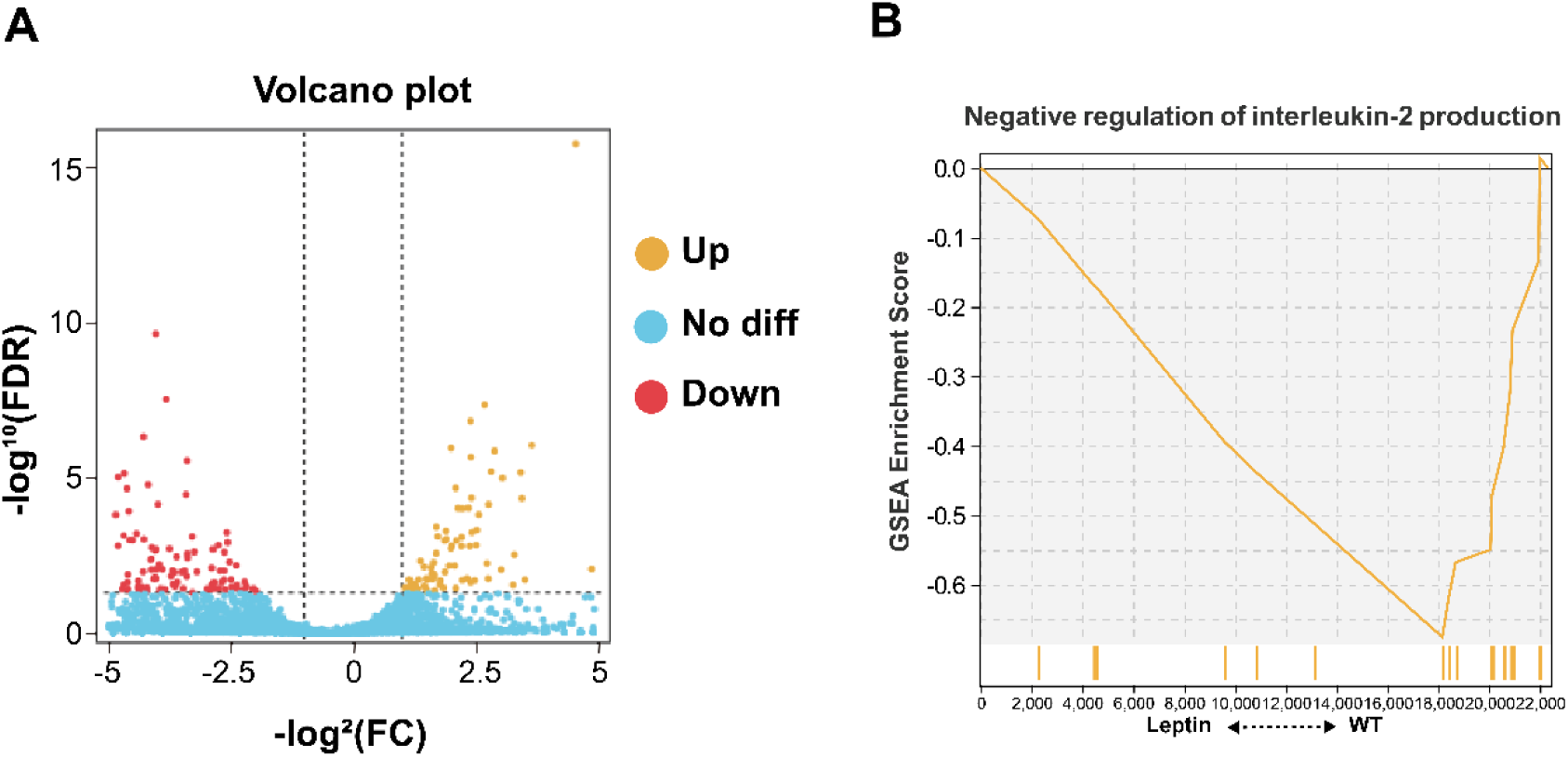
**(A)** Volcano plots of DEGs in leptin and WT group **(B)** The GSEA result of ovarian inflammation.

**Fig. S2:**
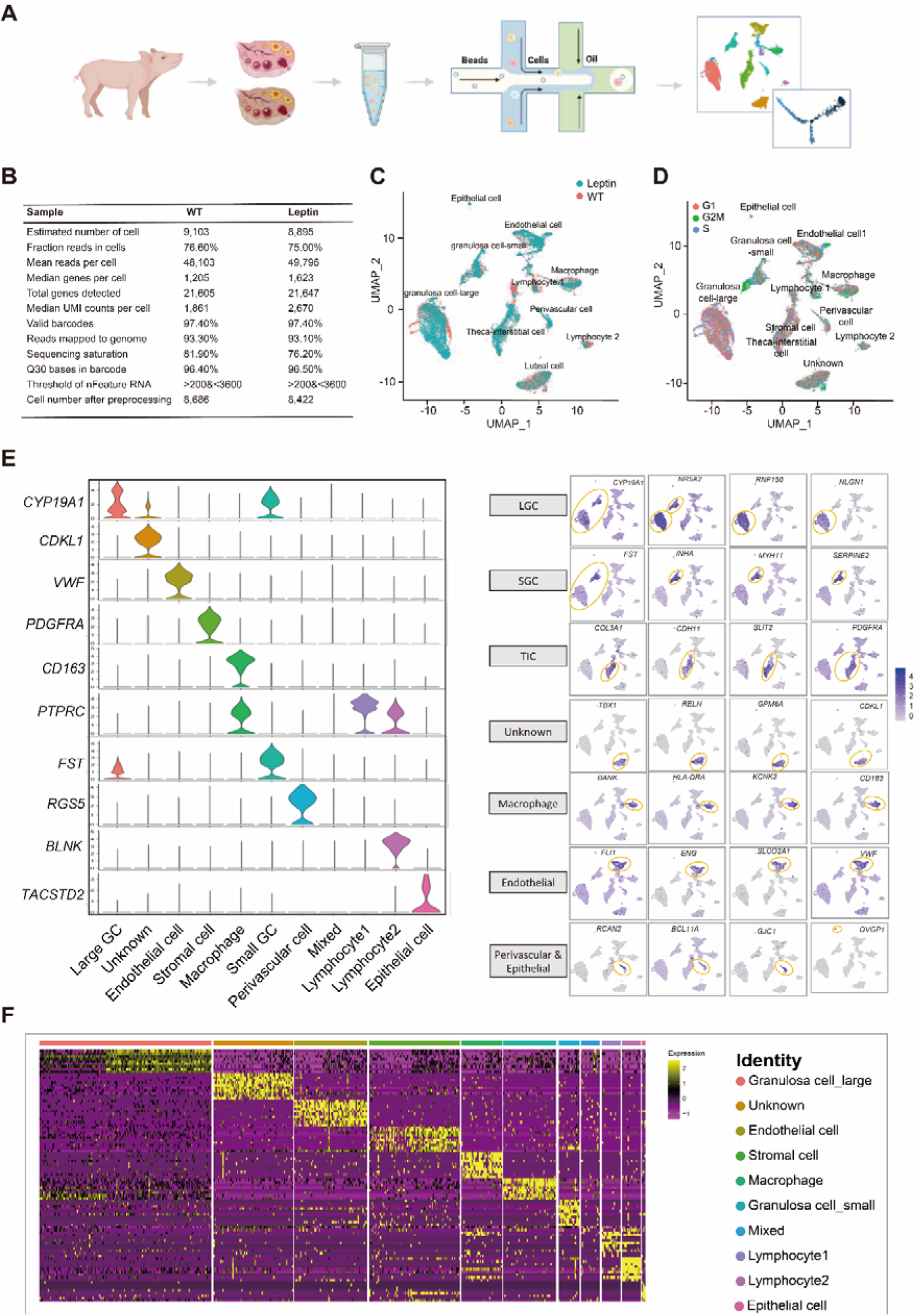
Sequencing information and novel marker genes. **(A)** Schematic diagram of snRNA sequencing. **(B)** Summary information of sample data identified by CellRanger. **(C)** UMAP plot showing sample identity (WT and leptin) **(D)** UMAP clustering of 3 phases in different cell populations. **(E)** Distribution of feature genes at different cell clusters. The gene expression levels are indicated by the colors of the bar. **(F)** Heatmap expression of upregulated genes in different cell populations.

**Fig. S3:**
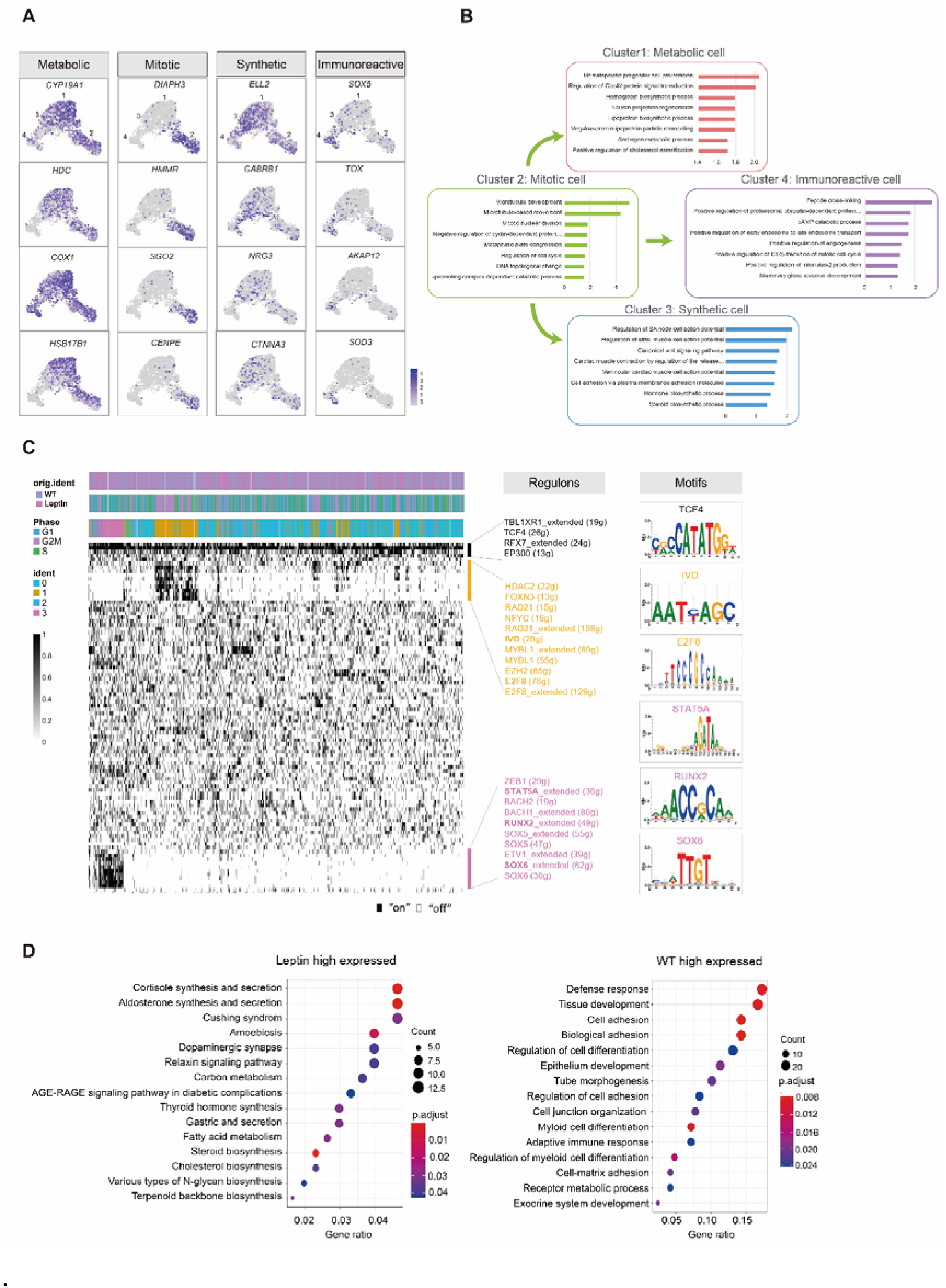
Gene expression pattern of ovarian cell subtypes. **(A)** Distribution of feature genes at four different clusters of SGC. The gene expression levels are indicated by the colors of the bar. **(B)** Representative GO terms of highly expressed genes in leptin and WT pig ovaries. **(C)** Mapping regulons activity of transcriptional factors in SGCs using SCENIC. **(D)** Representative KEGG terms of highly expressed genes in leptin group (left) and GO Biological Process terms of highly expressed genes in WT group (right).

**Table S1:**
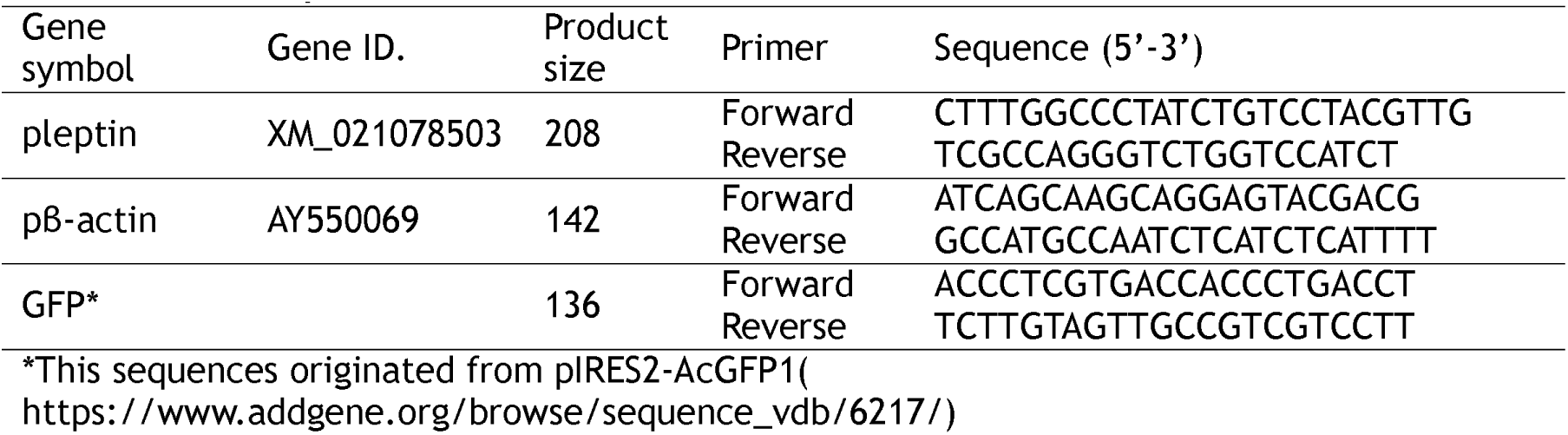
List of primers for PCR.

**Table S2:**
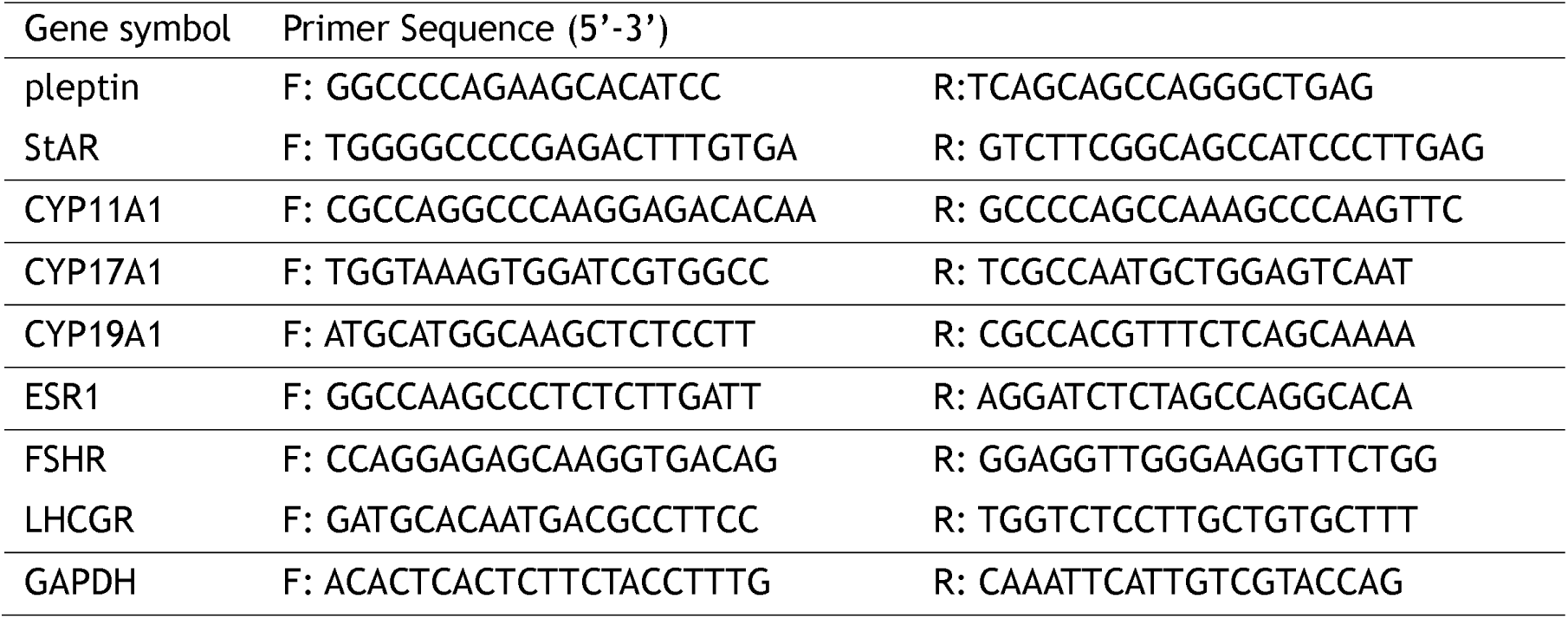
List of primers for qPCR.

**Table S3:** List of proteins.

| **Protein Name** | **Cas#** | **Company** | **Source** | **Dose** |
| --- | --- | --- | --- | --- |
| anti-β-actin (β-actin) | A5441 | Sigma-Aldrich | Mouse | 1:2000 |
| FSHR Polyclonal antibody | 22665-1-AP | Proteintech | Rabbit | 1:2000 |
| LHCGR Polyclonal antibody | 19968-1-AP |  |  | 1:1000 |
| StAR Polyclonal antibody | 12225-1-AP |  |  | 1:1000 |
| Cytochrome P450 17A1 Antibody (CYP17A1) | 14447-1-AP |  |  | 1:1000 |
| Cytochrome P450 19A1 Antibody (CYP19A1) | AF5229 | Affinity | Rabbit | 1:2000 |
| ERα (F-10) (ESR1) | SC-8002 | Santa Cruz Biotechnology, Inc | Mouse | 1:1000 |
| Caspase-3 Antibody | #9662 | Cell Signaling Technology | Rabbit | 1:1000 |

